# Soft robotics-enabled large animal model of HFpEF hemodynamics for device testing

**DOI:** 10.1101/2023.07.26.550654

**Authors:** Luca Rosalia, Caglar Ozturk, Sophie X. Wang, Diego Quevedo-Moreno, Mossab Y. Saeed, Adam Mauskapf, Ellen T. Roche

## Abstract

Heart failure with preserved ejection fraction (HFpEF) is a major challenge in cardiovascular medicine, accounting for approximately 50% of all cases of heart failure. Due to the lack of effective therapies for this condition, the mortality associated with HFpEF remains higher than that of most cancers. Despite the ongoing efforts, no medical device has yet received FDA approval. This is largely due to the lack of an in vivo model of the HFpEF hemodynamics, resulting in the inability to evaluate device effectiveness in vivo prior to clinical trials. Here, we describe the development of a highly tunable porcine model of HFpEF hemodynamics using implantable soft robotic sleeves, where controlled actuation of a left ventricular and an aortic sleeve can recapitulate changes in ventricular compliance and afterload associated with a broad spectrum of HFpEF hemodynamic phenotypes. We demonstrate the feasibility of the proposed model in preclinical testing by evaluating the hemodynamic response of the model post-implantation of an interatrial shunt device, which was found to be consistent with findings from in silico studies and clinical trials. This work addresses several of the limitations associated with previous models of HFpEF, such as their limited hemodynamic fidelity, elevated costs, lengthy development time, and low throughput. By showcasing exceptional versatility and tunability, the proposed platform has the potential to revolutionize the current approach for HFpEF device development and selection, with the goal of improving the quality of life for the 32 million people affected by HFpEF worldwide.

## INTRODUCTION

Heart failure with preserved ejection fraction (HFpEF) is a clinical syndrome in which patients have signs and symptoms of heart failure despite a normal left ventricular ejection fraction (LVEF; ≥ 50%) (*1*). HFpEF accounts for approximately 50% of all patients with heart failure, resulting in over 3 million people with this condition in the US and up to 32 million globally (*2, 3*). Currently, pharmacological therapies have shown limited mortality benefits and no medical device has yet been approved by the FDA for the treatment of HFpEF. As a result, the mortality associated with HFpEF is higher than that of most cancers (*4, 5*).

HFpEF encompasses conditions with diverse etiologies, usually with multisystem involvement, entailing cardiac, pulmonary, vascular, metabolic, renal, and/or hepatic abnormalities (*6, 7*). Our understanding of the pathophysiological processes that lead to these changes remains incomplete and hypotheses are constantly evolving. From a biomechanical standpoint, HFpEF is characterized by left ventricular (LV) stiffening or loss of compliance, hindering its ability to relax during diastole and reducing its capacity to fill (*8*). Loss of LV compliance can result from a variety of conditions, including metabolic disease, microvascular inflammation, restrictive, infiltrative, and hypertrophic cardiomyopathies, or can be secondary to pressure overload (or increased afterload) due to hypertension or aortic stenosis (*9–15*). In the context of pressure overload, concentric remodeling or thickening of the LV is thought to ensue as a compensatory mechanism to minimize changes in LV wall stress, governed by the law of Laplace (*16*). Such loss of LV compliance causes an upward shift of the end-diastolic pressure-volume relationship (EDPVR), which results in a drop in the LV filling volumes, or end-diastolic volume (EDV), and higher filling pressures or LV end-diastolic pressure (LVEDP) (*17*). As a result, the amount of blood pumped during each heartbeat, or stroke volume (SV), is diminished, driving a drop in cardiac output (CO) and the inability of the heart to meet the metabolic demands of the body. Evidence of increased filling pressures (LVEDP, pulmonary capillary wedge pressure, or right atrial pressure) at rest or with exercise is required for a clinical diagnosis of HFpEF (*18*).

Elevated LV filling pressures are transmitted retrogradely to the left atrium (LA) and the pulmonary circulation, resulting in a multitude of manifestations and complications of HFpEF. First, higher atrial pressures drive LA remodeling processes, which cause atrial fibrillation in 40-60% of the patients, increasing the risk of embolic stroke and rhythm abnormalities remarkably (*19*). Second, high pressures are transmitted to the pulmonary circulation, causing pulmonary hypertension (up to 80%) as well as symptoms of pulmonary edema and dyspnea on exertion, and eventually to the right heart, leading to right ventricular dysfunction (up to 50%) and other complications (*20, 21*). Further, symptom management is often complicated by autonomic imbalance, characterized by the upregulation of sympathetic activity with withdrawal of the vagal tone, further increasing the risk for atrial fibrillation and chronotropic incompetence (*22, 23*).

First-line treatment for HFpEF consists of sodium-glucose cotransporter type 2 inhibitors, which reduce hospitalization and cardiovascular death by approximately 20%, although the primary mechanism underlying their clinical benefits remains unclear (*24, 25*). Loop diuretics are also prescribed in patients with overt congestion and aerobic exercise is broadly recommended (*2, 26*). In addition, clinical management aims to address any known underlying causes or complications of HFpEF (*27*). In tandem with the search for pharmacologic therapies that can improve the survival and quality of life of HFpEF patients, several of which are currently undergoing clinical trials, substantial efforts have been made toward the development of device-based solutions (*28, 29*). Medical devices for HFpEF seek to restore adequate cardiac mechanics or hemodynamics through a variety of mechanisms. The primary classes of devices for HFpEF include atrial shunts, LV expanders, and mechanical circulatory support (MCS) devices.

Atrial shunts are designed to lower elevated LA pressures by shunting blood from the LA to lower-pressure structures, such as the right atrium (RA) or the coronary sinus. LA-to-RA interatrial shunts include the interatrial shunt device (IASD, Corvia Medical Inc), the V-Wave shunt (V-Wave Ltd), and the atrial flow regulator (AFR) device (Occlutech). These devices are made of a Nitinol frame or mesh of various geometries that creates a 6-10 mm opening between the LA and RA to shunt blood down its pressure gradient (*30–32*). Recently, studies have investigated the feasibility of device-free interatrial shunting (*33*). An alternative design by Edwards LifeSciences (Transcatheter Atrial Shunt system) aims to prevent RA overload by creating a 7-mm shunt between the LA and the coronary sinus (*34*). Another class of devices is that of LV expanders, which store mechanical energy during contraction – transferring it to the LV during diastole – seeking to augment LV filling capacity. Two main types of LV expanders have been designed to date, namely the ImCardia (CorAssist Inc.) device, which is an extracardiac expander designed for implantation on the epicardial surface, now withdrawn due to safety concerns, and the endocardial CORolla (CorAssist Inc.) device for transapical implantation (*35*). Finally, MCS strategies involve pumps to reduce LA pressures and increase CO (*36*). Examples of MCS devices include the PulseVad by Northern Development, which is a minimally invasive smart pump aiming to enable adaptive flow from the LA to the descending aorta, the left atrial assist device (LAAD) – a continuous-flow LA-to-LV pump designed to be implanted at the level of the mitral valve, and the CoPulse device – a type of valveless pulsatile pneumatic pump made of a flexible polyurethane membrane that is designed for apical implantation (*37, 38*). In addition, studies have been conducted to investigate the feasibility of traditional LVAD devices for use in HFpEF patients, although concerns remain over the risk of LA suction or inflow cannula obstruction due to concentric LV remodeling (*39*).

These device-based solutions for HFpEF are all at different stages of development, with interatrial shunts having been recently marketed for use in the European Union, the CORolla LV expander undergoing clinical trials, and MCS devices in various preclinical testing phases. However, to date, none of these devices have received FDA approval (*29*). A major limitation to the development of these devices and their market approval in the US is the lack of an adequate in vivo model of the HFpEF hemodynamics (*40, 41*). As a result, preclinical safety and functional evaluation of device-based solutions for HFpEF can only be conducted on healthy animals, considerably hindering their reliability (*35, 42*).

Current large animal models of HFpEF suffer from a variety of limitations, including their limited hemodynamic fidelity, elevated costs, lengthy development times, and low throughput. These models can only rarely successfully re-create the elevations in filling pressures associated with HFpEF, require follow-ups ranging between 6-24 weeks, and are affected by mortality rates of 30% or higher (*41*). Most of these models aim to recapitulate the structural and functional changes associated with the disease by inducing pressure overload by means of an aortic cuff, band, or stent (*43–45*). Another method to induce pressure overload involves inducing renovascular hypertension via renal wrapping, renal clamping, renal embolization, or via the administration of deoxy-corticosterone acetate (DOCA) salt (*46–48*). High-fat diets are often leveraged as a supplemental method for these models, aiming to recapitulate changes in metabolic function associated with HFpEF (*49*). Another method currently under development involves reducing LV filling by the use of an intraventricular balloon; however, this technique is still lacking a post-operative functional evaluation and may induce systolic dysfunction, which is only rarely associated with HFpEF (*41*). Because of all these limitations, no such models are currently being used for the evaluation of medical devices for HFpEF.

In this work, we describe the development of a large animal model of HFpEF hemodynamics that can recapitulate acute changes in LV biomechanics and afterload associated with disease in a tunable and controllable manner. The model is enabled by two implantable soft robotic sleeves; an LV sleeve that re-creates the loss of LV compliance characteristic of HFpEF, and an aortic sleeve that can recapitulate changes in the afterload associated with most hemodynamic phenotypes of HFpEF (Fig. 1A). In previous studies, we demonstrated the ability of the LV sleeve to limit cardiac filling in an in vitro model of aortic stenosis and that of the aortic sleeve to recreate the hemodynamics of pressure overload in vitro and in vivo (*50, 51*). Here, we redesign the LV sleeve for the development of the first porcine model of HFpEF capable of tuning pressure overload and ventricular compliance independently, in a facile, immediate, tunable manner, which allows us to re-create a broad spectrum of HFpEF hemodynamics. By investigating the hemodynamic response of the model post-implantation of an interatrial shunt device, we then demonstrate the applicability of this model for in-vivo evaluation of device-based solutions for HFpEF (Fig. 1B). We believe that this proposed porcine model of HFpEF hemodynamics can establish a new standard for the development of device-based solutions for HFpEF, addressing an unmet need in cardiovascular medicine by supporting translational efforts towards the development of more effective treatment strategies for people with HFpEF.

**Fig. 1.**
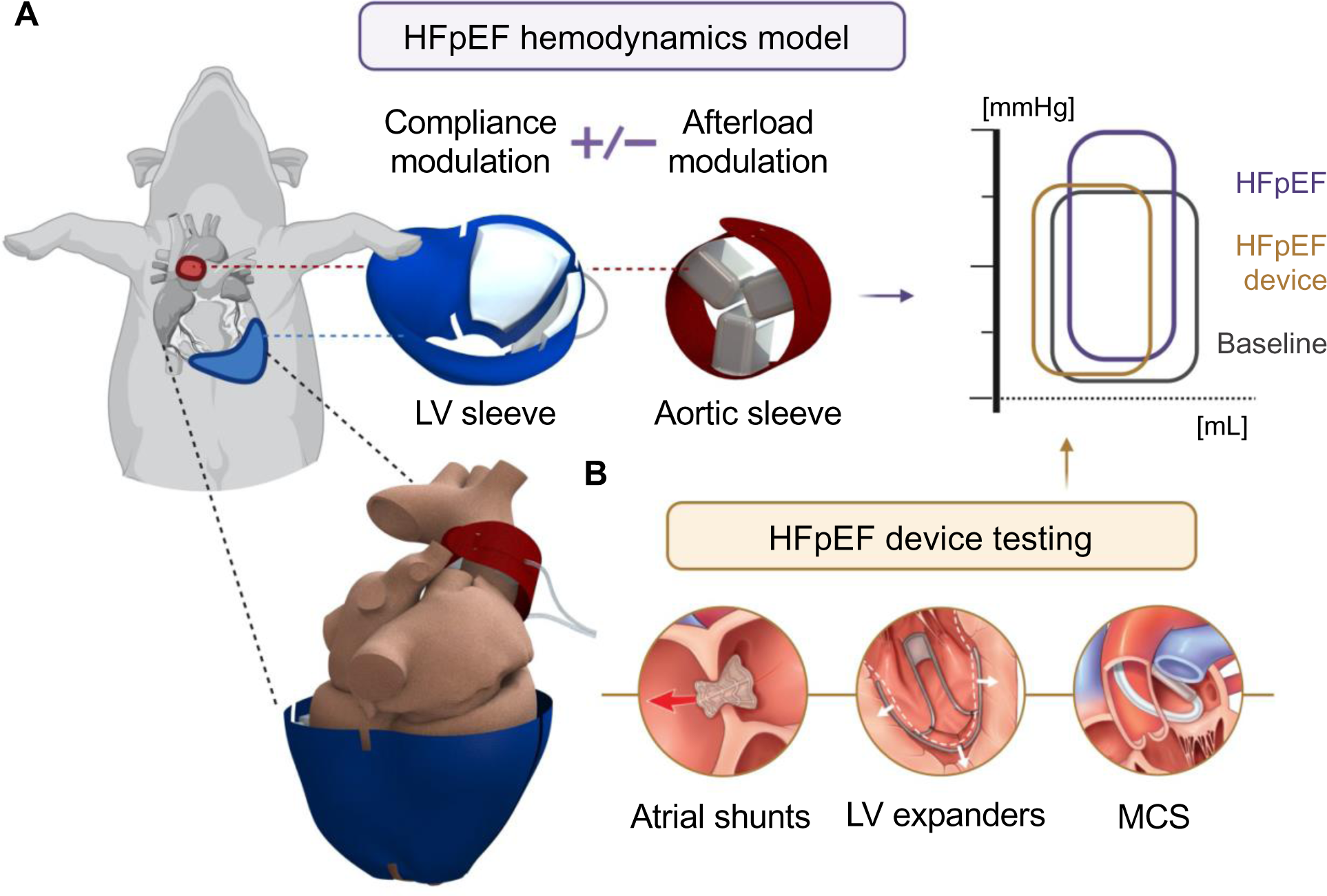
Overview of the acute porcine model of heart failure with preserved ejection fraction (HFpEF). **(A)** Model of the HFpEF hemodynamics enabled by implantable soft robotic sleeves. Actuation of the left ventricular (LV) sleeve causes reduced LV compliance, while afterload modulation can be achieved using an aortic sleeve. **(B)** Applicability of the model for in vivo testing of device-based solutions for HFpEF. MCS = mechanical circulatory support.

## RESULTS

### Left ventricular sleeve can modulate cardiac compliance to re-create HFpEF hemodynamics in a porcine model

We designed an LV sleeve that can be implanted around the epicardial surface of the heart in a swine model to modulate cardiac filling function. The sleeve (Fig. 2A) is composed of two inflatable pockets made of thermoplastic polyurethane (TPU, HTM 8001-M 80A shore polyether film, 0.012 inches in thickness; American Polyfilm Inc.) that expand under pressure and of a 200-Denier inelastic TPU-coated fabric (Oxford fabric, Seattle fabrics Inc.) that directs the expansion of the TPU pockets towards the LV and enables anchoring of the sleeve onto the epicardial surface. The sleeve is designed to be implanted so that the inflatable pockets are in contact with the outer wall of the LV, whereas the rest of the fabric wraps around the right ventricle (RV). Velcro straps were used to secure the fabric tightly around the epicardial wall. The TPU pockets were designed to conform with the LV anatomy; they consist of a double layer of TPU sheets that are first vacuum formed to 3D printed (Veroblue, Objet 30 3D printer, Stratasys) molds, and then heat-sealed at their edges to create a fully enclosed compartment. The pockets were then heat-sealed to the fabric and connected to their respective actuation line (latex rubber 1/16-inch inner diameter and 1/8-inch outer diameter tubing; McMaster-Carr). Actuation of the LV sleeve results in a drop in LV compliance indicated as an upward shift of the end-diastolic pressure-volume relationship (EDPVR) in the LV pressure-volume (PV) loop (Fig. 2A).

**Fig. 2.**
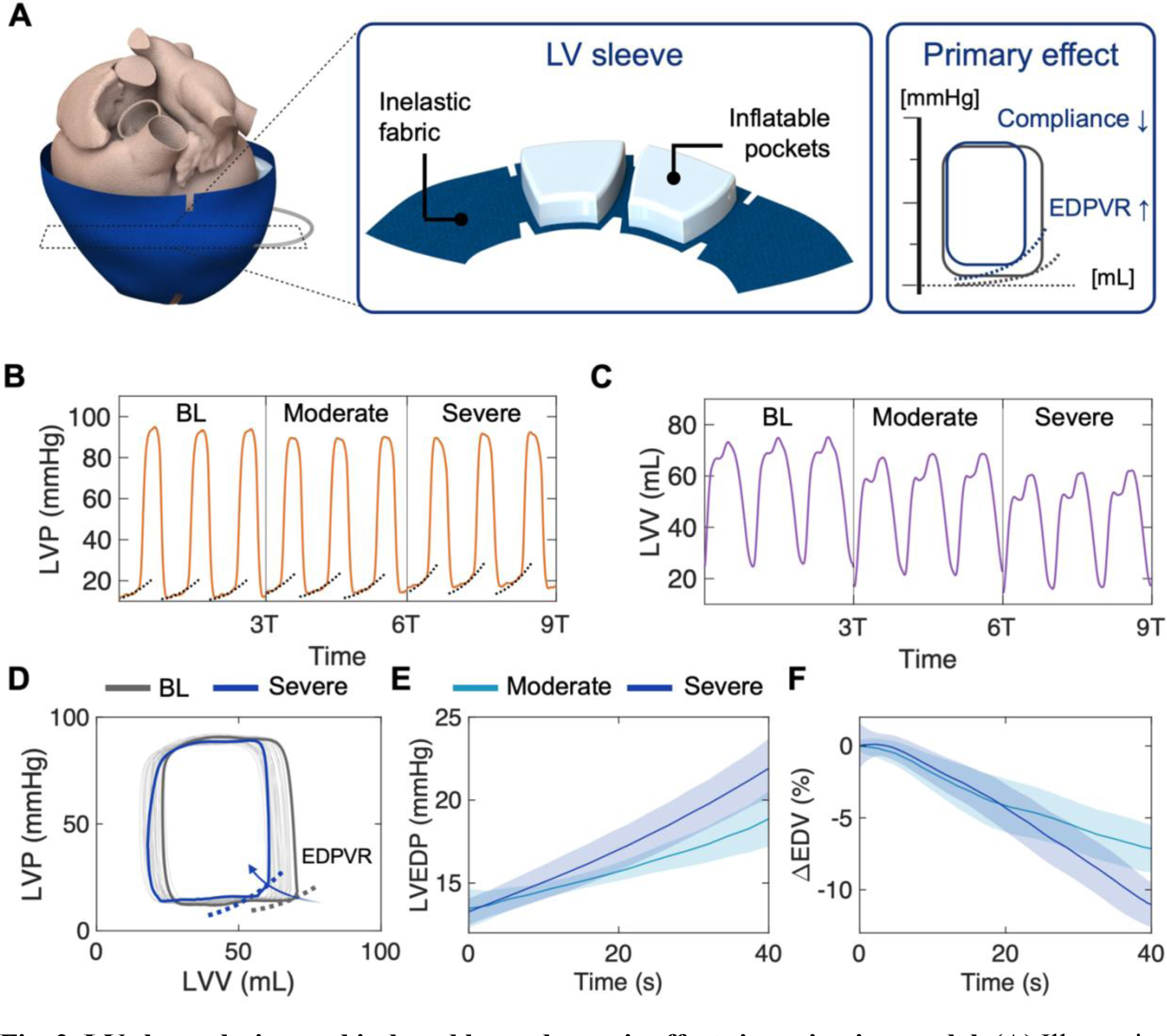
LV sleeve design and induced hemodynamic effects in an in vivo model. **(A)** Illustration of the LV sleeve, with details of the inelastic fabric and the inflatable pockets. Changes in the LV PV loop, illustrating a drop in LV compliance as the primary hemodynamic effect induced by actuation of the LV sleeve. **(B-C)** Representative **(B)** LVP and **(C)** LVV waveforms for three consecutive heartbeats at baseline (BL) and for moderate and severe actuation. The dotted lines in **(B)** highlight elevations in filling pressures. **(D)** Representative PV loop progression from BL to severe actuation, highlighting an increase in EDPVR. **(E-F)** LVEDP and changes in EDV during LV sleeve actuation. Data show mean ± 1 S.D. Each test was conducted on n = 3 swine and repeated n = 3 times. EDPVR = end-diastolic pressure-volume relationship; LVP = left ventricular pressure; LVV = left ventricular volume; T = heart cycle period; LVEDP = left ventricular end-diastolic pressure; EDV = end-diastolic volume.

In a total of n = 3 swine, the LV sleeve was actuated at various volumes, using a syringe pump over a period of approximately 40 seconds (see Materials and Methods section). Fig. 2 B-C show representative changes in LV pressures (LVP) and volumes (LVV), respectively, achieved at moderate and severe actuation compared to baseline. The dotted lines in Fig. 2B highlight a progressive increase in the filling pressures, whereas Fig. 2C shows a corresponding drop in the filling volumes or maximum LVV. The progression of the LV PV loop from baseline to severe actuation corroborates these changes and confirms the expected upward shift of the EDPVR, as an indication of limited LV filling induced by actuation of the LV sleeve (Fig. 2D). Fig. 2 E-F illustrates the evolution of these hemodynamic changes averaged across the n = 3 trials, during moderate and severe actuation of the LV sleeve. From a baseline value of LVEDP_BL_ = 13.3 ± 1.2 mmHg, the LVEDP increased to 18.9 ± 1.7 mmHg and 21.9 ± 1.8 mmHg at moderate and severe actuation, respectively. Correspondingly, actuation caused a drop in the EDV equal to ΔEDV_M_ = -7.1 ± 1.6 % and ΔEDV_S_ = -11.0 ± 1.6 %. Overall, actuation of the LV sleeve alone was shown to recapitulate changes in LV compliance, which is the primary hemodynamic hallmark of HFpEF.

### Aortic sleeve can recapitulate pressure overload driving HFpEF

Being able to induce changes in the afterload in conjunction with LV filling modulation is critical to recapitulate a broader spectrum of HFpEF hemodynamic phenotypes. To this end, we integrated our in vivo model of HFpEF with a soft robotic aortic sleeve that can be controlled to re-create elevations in afterload often responsible for or associated with HFpEF. Here, we describe the changes in the LV and aortic hemodynamics induced by the aortic sleeve only, before characterizing the hemodynamics resulting from combining both sleeves in the following section. The aortic sleeve is made of three inflatable TPU pockets that expand under pressure to compress the ascending aorta, inducing pressure overload in a controllable fashion (Fig. 3A). Analogous to the LV sleeve, the three pockets are connected to an actuation line and to an inelastic fabric layer that directs the expansion of the pockets inwards and secures the sleeve around the aortic anatomy (Fig. 3A). The materials and manufacturing workflow, involving vacuum forming and heat sealing, of the aortic sleeve are the same as those of the LV sleeve. The aortic sleeve was actuated using the same syringe pump over a period of 5 seconds at two distinct actuation levels, namely moderate and severe actuation (see Materials and Methods section).

**Fig. 3.**
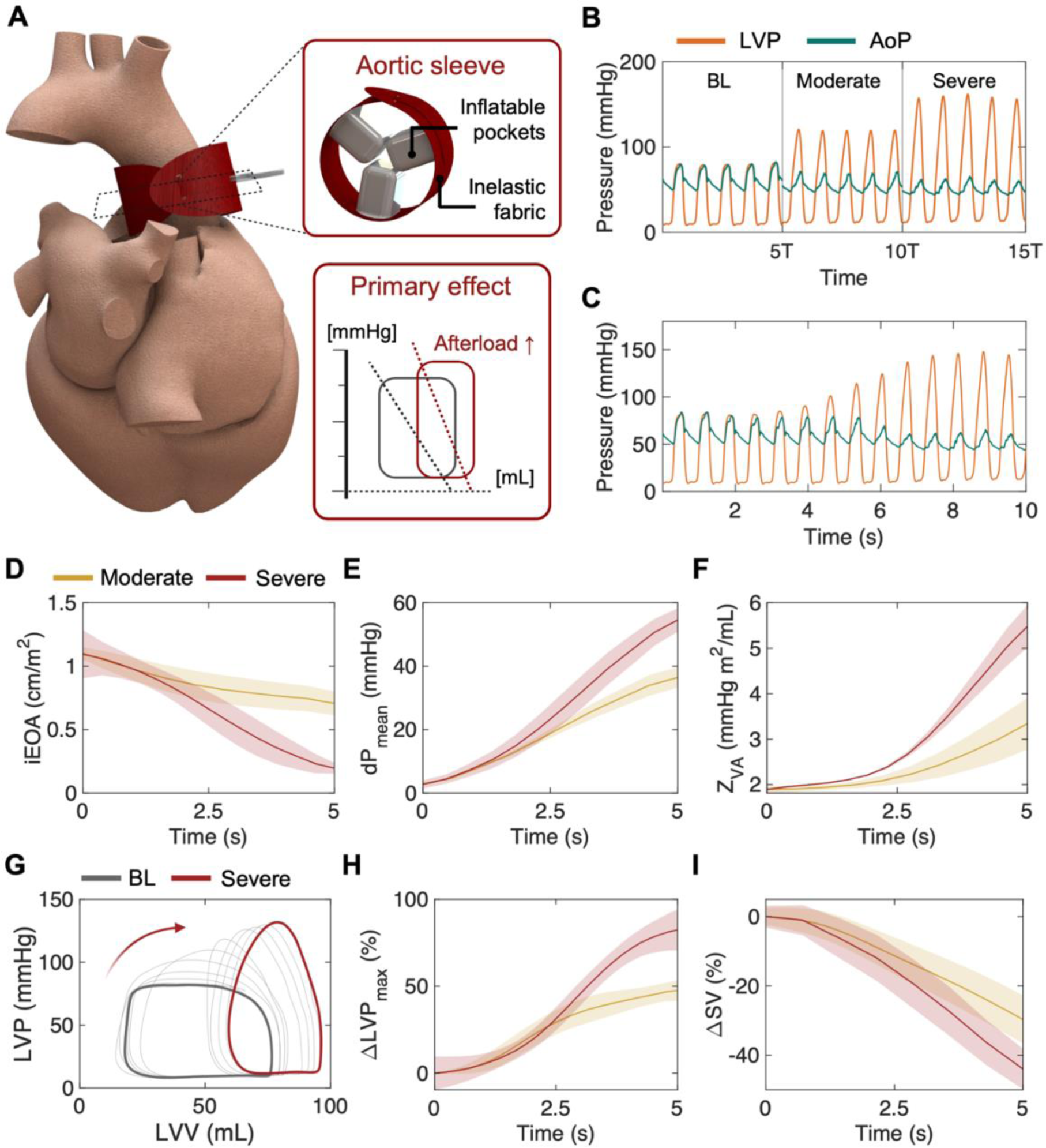
Hemodynamic effects of the soft robotic aortic sleeve for in vivo model of HFpEF. **(A)** Overview of the soft robotic aortic sleeve, illustrating the inflatable pockets and the inelastic fabric layer. Exemplary PV loop highlights elevations in afterload from baseline as induced by the aortic sleeve. **(B-C)** Representative LVP and AoP waveforms **(B)** at discrete levels, including (BL) baseline, moderate and severe actuation for five consecutive heartbeats, and **(C)** progressively during actuation. **(D-F)** Changes in aortic hemodynamic metrics due to moderate and severe actuation of the aortic sleeve, including **(D)** iEOA, **(E)** dP_mean_, and **(F)** Z_VA_. **(G)** Representative evolution of PV loops from BL due to severe aortic sleeve actuation. **(H-I)** Changes in **(H)** LVP and **(I)** SV for moderate and severe actuation. Data show mean ± 1 S.D. Each test was conducted on n = 3 swine and repeated n = 3 times. LVP = left ventricular pressure; AoP = aortic pressure; T = heart cycle period; iEOA = indexed effective orifice area; dP = transaortic pressure gradient; Z_VA_ = valvulo-arterial impedance; LVV = left ventricular volume; SV = stroke volume.

Fig. 3 B-C shows representative changes in LVP and aortic pressure (AoP) waveforms induced by actuation of the aortic sleeve. Fig. 3B highlights these changes from baseline to moderate and severe actuation in a discrete fashion, whereas the evolution of the waveforms over time is visualized in Fig. 3C. Consistently, these graphs show that actuation of the aortic sleeve leads to elevations in the systolic LVP and a reduction of the AoP, establishing a pressure gradient between the two. Evaluation of hemodynamic parameters used for clinical assessment of aortic stenosis showed a progressive drop in the indexed effective orifice area (iEOA_BL_ = 1.09 ± 0.19 cm/m^2^, iEOA_M_ = 0.71 ± 0.09 cm/m^2^, iEOA_S_ = 0.20 ± 0.05 cm/m^2^; Fig. 3D) caused by controlled aortic constriction. This led to an increase in the mean transaortic pressure gradient (dP_mean_BL_ = 2.7 ± 1.3 mmHg, dP_mean_M_ = 36.4 ± 3.2 mmHg, dP_mean_S_ = 54.6 ± 3.7 mmHg; Fig. 3E) and in elevations in the valvulo-arterial impedance (Z_VA_BL_ = 1.9 ± 0.1 mmHg m^2^/mL, Z_VA_M_ = 3.3 ± 0.6 mmHg m^2^/mL, Z_VA_S_ = 5.5 ± 0.5 mmHg m^2^/mL; Fig. 3F).

Representative changes in cardiac function during actuation of the aortic sleeve are shown in Fig. 3 G-I. As predicted, controlled aortic constriction led to a rightward shift of the LV PV loop, with an increase in systolic LVP and a drop in the SV (Fig. 3G). Quantitatively, moderate and severe actuation led to changes in LVP and SV equal to ΔLVP_max_M_ = 47.6 ± 6.0 % and ΔLVP_max_S_ = 82.4 ± 11.8 % (Fig. 3H) and ΔSV_M_ = -29.7 ± 7.0 % and ΔSV_S_ = -43.9 ± 5.9 % (Fig. 3I).

### Soft robotic sleeves can be modulated to recapitulate a spectrum of HFpEF hemodynamics

Through the combination of the LV and aortic sleeves, our porcine model of HFpEF can re-create various hemodynamic phenotypes of HFpEF, which will allow us to capture a broad spectrum of HFpEF etiologies and disease severities. The illustration in Fig. 4A shows representative combinations of the LV and aortic sleeve actuation schemes that can be leveraged to recapitulate various HFpEF hemodynamics. When the aortic sleeve is off, actuation of the LV sleeve can capture the hemodynamics of limited LV filling associated with HFpEF in the absence of pressure overload, for example, relevant for patients with restrictive, infiltrative, or hypertrophic cardiomyopathies. Conversely, actuation of both sleeves enables afterload and LV compliance modulation for the recapitulation of the hemodynamics of HFpEF caused by (or comorbid with) aortic stenosis or hypertension. The schematic in Fig. 4A also shows that the actuation levels of the sleeves can be modulated individually to recreate various severities of disease.

**Fig. 4.**
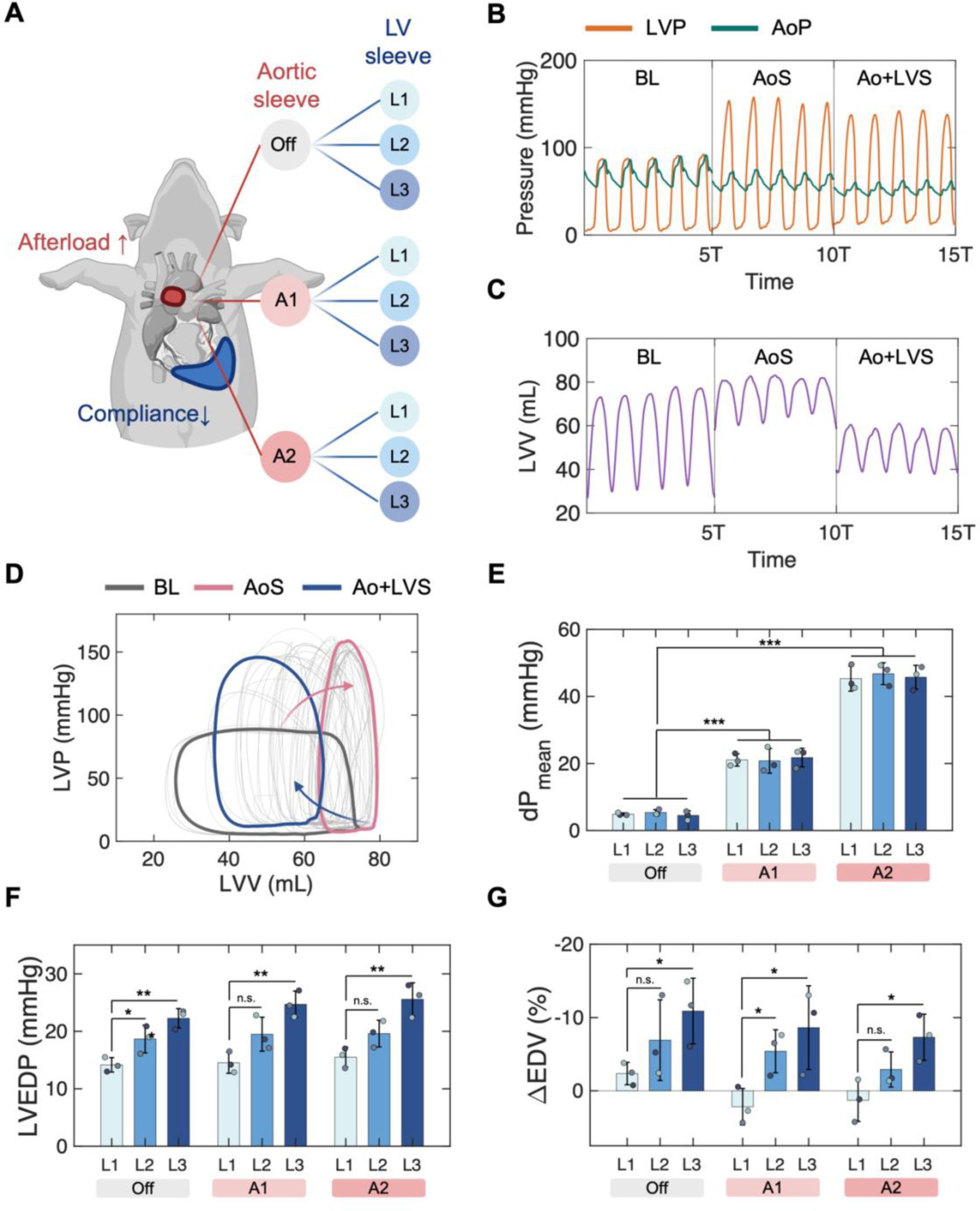
In vivo HFpEF hemodynamic modulation via integration of left ventricular (LV) and aortic sleeves. **(A)** Schematic illustrating representative actuation patterns of LV and aortic sleeves. **(B-C)** Representative **(B)** LVP and AoP and **(C)** LVV waveforms for five consecutive heartbeats at baseline (BL) and during actuation of the aortic sleeve (AoS) and of both aortic and left ventricular sleeves (Ao+LVS). **(D)** Representative LV PV loop progression from baseline through pressure overload (AoS) to pressure overload and HFpEF (Ao+LVS). **(E-G)** Metrics of pressure overload and HFpEF, including **(E)** dP_mean_, **(F)** LVEDP, and **(G)** EDV for various actuation schemes of the aortic (bottom: off, A1, A2) and LV (top: L1-L3) sleeves. Data show mean ± 1 S.D. Each test was conducted on n = 3 swine and repeated n = 3 times. *: *P* < 0.05, **: *P* < 0.01, ***: *P* < 0.001, n.s.: non-significant. LVP = left ventricular pressure; AoP = aortic pressure; LVV = left ventricular volume; T = heart cycle period; dP = transaortic pressure gradient; LVEDP = left ventricular end-diastolic pressure; EDV = end-diastolic volume; L1-3 = actuation levels.

Fig. 4 B-D illustrates hemodynamic changes induced by actuation of the aortic sleeve only (AoS), then followed by actuation of both sleeves (Ao+LVS). This allowed us to investigate the ability of our model to simulate the hemodynamic progression of patients with HFpEF secondary to pressure overload in a longitudinal manner. Fig. 4 B-C shows that actuation of the LV following actuation of the aortic sleeve can cause an increase in filling pressures and a drop in filling volume, even in the context of pressure overload. Further, Fig. 4B highlights that the systolic LVP may drop slightly due to actuation of the LV sleeve, which is a consequence of the decrease in filling volumes (or preload). Similarly, actuation of the LV sleeve induces a considerable drop in both the EDV and the end-systolic volume (ESV), as seen in Fig. 4C. Overall, the SV is reduced in Ao+LVS compared to baseline, which is consistent with the effects of limited cardiac filling associated with HFpEF. These changes can be visualized in Fig. 4D, which shows the progression of PV loops from baseline to pressure overload only (AoS), and finally to both pressure overload and limited cardiac filling (Ao+LVS).

The graphs in Fig. 4 E-G showcase the broad spectrum of hemodynamic characteristics that could be obtained by selective and independent actuation of the LV and aortic sleeves. Changes in dP_mean_ in Fig. 4E show that the degree of pressure overload can be reliably controlled by the aortic sleeve (dP_mean_Off_L1_ = 4.9 ± 0.4 mmHg, dP_mean_A1L1_ = 21.1 ± 1.8 mmHg, dP_mean_A2L1_ = 45.3 ± 3.7 mmHg, *P* < 0.001; Fig. 4E) and that progressively reducing filling capacity through actuation of the LV sleeve does not cause significant alterations to it (dP_mean_A2L3_ = 45.7 ± 3.6 mmHg; Fig. 4E). Actuation of the LV sleeve resulted in an increase in the LVEDP (LVEDP_Off_L1_ = 14.2 ± 1.3 mmHg, LVEDP_Off_L3_ = 22.3 ± 1.7 mmHg, *P* < 0.01; Fig. 4F). When combined with pressure overload, and for the same actuation level of the LV sleeve, these changes become slightly more pronounced (LVEDP_A2L1_ = 15.5 ± 1.7 mmHg, LVEDP_A2L3_ = 25.6 ± 2.9 mmHg, *P* < 0.01; Fig. 4F).

Similarly, a reduction in end-diastolic volumes ranging from 2-10% can be achieved, depending on the level of actuation (ΔEDV_Off_L1_ = -2.4 ± 1.5 %, ΔEDV_Off_L3_ = -10.9 ± 4.5 %, *P* < 0.05; Fig. 4F). Because pressure overload is also associated with an increase in EDV (Fig. 4F) when the LV sleeve is used in tandem with the aortic sleeve, its actuation level can be adjusted to obtain a desired change in EDV (ΔEDV_OFF_L1_ = -2.4 ± 1.5 %, ΔEDV_A2L2_ = -2.9 ± 3.2 %; Fig. 4F). These data demonstrate that by varying the actuation levels independently for each sleeve, this model can re-create a variety of hemodynamic profiles of relevance for studies of HFpEF and device testing.

### In vivo HFpEF hemodynamic model enables high-fidelity testing of device-based solutions

Fine control over the hemodynamic characteristics of HFpEF in a preclinical model enables high-fidelity testing of medical devices for this condition. To demonstrate feasibility in device testing, we simulated HFpEF intervention by creating a device-free atrial septal defect (ASD) and by implantation of an interatrial shunt device (Fig. 5A). Details of the interventional approaches can be found in the Materials and Methods section. Fluoroscopy and color Doppler imaging confirmed the correct positioning and patency of the interatrial shunt device (Fig. 5B).

**Fig. 5.**
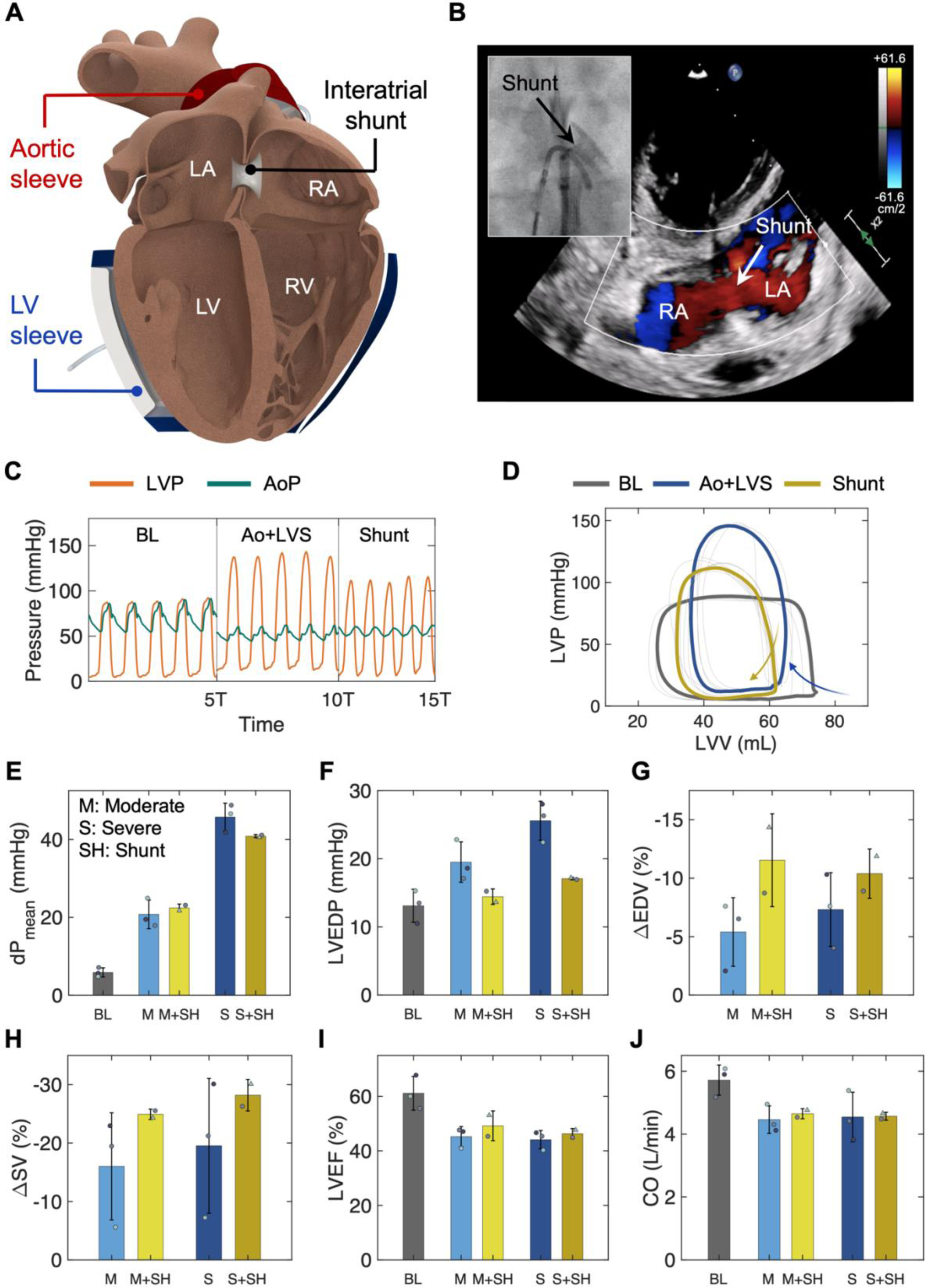
Application of HFpEF hemodynamic model for device testing and evaluation. **(A)** Illustration of aortic and LV sleeves and of an interatrial shunt used to simulate intervention in HFpEF in vivo model. **(B)** Color flow map imaging illustrating shunt patency and interatrial flow. Detail of shunt positioning on fluoroscopy. **(C-D)** Representative **(C)** LVP and AoP waveforms for five consecutive heartbeats and **(D)** LV PV loops at baseline (BL) and during actuation of the aortic (AoS) and of both sleeves (Ao+LVS). **(E-J)** Metrics of aortic and LV hemodynamics at BL, for moderate actuation (M), moderate actuation with shunt (M+SH), severe actuation (S), and severe actuation with shunt (S+SH), including **(E)** dP_mean_, **(F)** LVEDP, **(G)** EDV, **(H)** SV, **(I)** LVEF, and **(J)** CO. Data show mean ± 1 S.D. Each test was conducted on n = 3 swine and repeated n = 3 times for the HFpEF disease model and n = 2 for shunt evaluation. Triangles represent data points obtained with an AFR device (Occlutech). LA = left atrium; RA = right atrium; LV = left ventricle; RV = right ventricle; LVP = left ventricular pressure; AoP = aortic pressure; T = heart cycle period; LVV = left ventricular volume; dP = transaortic pressure gradient; LVEDP = left ventricular end-diastolic pressure; EDV = end-diastolic pressure; SV = stroke volume; LVEF = left ventricular ejection fraction; CO = cardiac output.

The hemodynamic effects of interatrial shunting on LV unloading can be visualized via the LVP and AoP waveforms and LV PV loops in Fig. 5 C-D. Both graphs show that shunting restores physiologic levels of filling pressures and that it causes a reduction in the systolic LVP. This occurs because interatrial shunting results in a drop in the preload, which causes the LV PV loop to move down its EDPVR curve, leading to a reduction in both peak systolic and end-diastolic pressures (Fig. 5D).

The drop in systolic LVP causes a decrease in the dP_mean_, which is more prominent when severe disease (both pressure overload and limited filling) is simulated (dP_mean_S_ = 45.7 ± 3.6 mmHg, dP_mean_S+SH_ = 40.8 ± 0.4 mmHg; Fig. 5E) compared to moderate disease (dP_mean_M_ = 20.8 ± 3.7 mmHg, dP_mean_M+SH_ = 22.4 ± 1.0 mmHg; Fig. 5E). LA-to-RA shunting restored levels of the LVEDP close to baseline (LVEDP_BL_ = 13.1 ± 2.4 mmHg, LVEDP_S_ = 25.6 ± 2.9 mmHg, LVEDP_S+SH_ = 17.1 ± 0.2 mmHg; Fig. 5F). Further, varying the level of disease severity was shown to have an effect on the LVEDP measured after intervention, where higher severity levels led to slightly elevated values of LVEDP (LVEDP_M+SH_ = 14.4 ± 1.1 mmHg, LVEDP_S+SH_ = 17.1 ± 0.2 mmHg; Fig. 5F). Analogously, the reduction in EDV induced by shunting was slightly more prominent for mild disease (ΔEDV_M_ = -5.4 ± 2.9 %, ΔEDV_M+SH_ = -11.5 ± 4.0 %; Fig. 5G) than for the simulated severe condition (ΔEDV_S_ = -7.3 ± 3.2 %, ΔEDV_S+SH_ = -10.4 ± 2.1 %; Fig. 5G).

Our model was able to predict changes in other metrics of cardiac function, including SV, LVEF, and CO, associated with intervention (Fig. 5 H-J). Due to a reduction in both the EDV and the ESV, we measured further drops in SV caused by shunting both when simulating mild (ΔSV_M_ = - 16.0 ± 9.2 %, ΔSV_M+SH_ = -25.9 ± 0.9 %; Fig. 5H) and severe (ΔSV_S_ = -19.5 ± 11.5 %, ΔSV_S+SH_ = -28.2 ± 2.7 %; Fig. 5H) HFpEF hemodynamics. This led to a slight increase in the LVEF due to shunting for both disease levels (LVEF_M_ = 45.2 ± 3.7 %, LVEF_M+SH_ = 49.2 ± 5.5 % LVEF_S_ = 44.1 ± 3.4 %, LVEF_S+SH_ = 46.3 ± 1.9 %; Fig. 5I). Finally, albeit modest, we observed an increase in CO (CO_M_ = 4.5 ± 0.4 L/min, CO_M+SH_ = 4.6 ± 0.2 L/min, CO_S_ = 4.5 ± 0.8 L/min, CO_S+SH_ = 4.6 ± 0.1 L/min; Fig. 5I), likely due to chronotropic compensation.

## DISCUSSION

This work presents the first large animal model of HFpEF hemodynamics for use in preclinical testing of medical devices for HFpEF. The proposed model is based on implantable soft robotic sleeves that can be controllably actuated to finely re-create changes in preload and afterload associated with (or leading to) HFpEF.

We describe the design, manufacturing, and development of a soft robotic LV sleeve to be implanted around the heart in a swine model to modulate LV compliance and filling capacity (Fig. 2A). The sleeve is made of pre-formed actuatable TPU pockets in contact with the epicardial surface of the LV. When pressurized, these pockets inflate to mechanically limit LV filling, leading to a controlled increase in LVEDP and a drop in the EDV and SV (Fig. 2 D-F), which represent the main hemodynamic signature of HFpEF. Elevations in the afterload (associated with up to 90% of cases of HFpEF) (*52*) were recapitulated using controlled aortic constriction using a soft robotic aortic sleeve (Fig. 3A). Analogous to the LV sleeve, the aortic sleeve is made of expandable TPU pockets attached to an inelastic fabric layer for implantation around the wall of the ascending aorta. By calculating diagnostic metrics of aortic stenosis (Fig. 3 D-F) and cardiac function (Fig. 3 G-I), we showed that actuation of the aortic sleeve can recapitulate various degrees of pressure overload that are clinically relevant for HFpEF.

In a swine model, we implanted both the LV and aortic sleeves and showed that they can be actuated in isolation or in combination to recapitulate a broad spectrum of HFpEF hemodynamic phenotypes (Fig. 4A). For each of the simulated conditions, we measured LVP, AoP (Fig. 4B), LVV (Fig. 4C), LV PV loops (Fig. 4D), dP_mean_ (Fig. 4E), LVEDP (Fig. 4F), and EDV (Fig. 4G) to investigate the effects of selective and progressive sleeve actuation on cardiac hemodynamics and confirmed that the changes in the hemodynamics obtained with our model are consistent with the clinical literature of HFpEF(*17*). We showed that actuation of the LV sleeve did not affect changes in the afterload induced by the aortic sleeve (Fig. 4E). Conversely, we observed an effect, albeit minor, of aortic sleeve actuation on the modulation of LVEDP and EDV induced by the LV sleeve. Particularly in the context of acute onset of pressure overload, severe elevations in the afterload may cause a rightward shift of the LV PV loop (Fig. 4D) characterized by an increase in the LVEDP and EDV. As a result, actuation of the aortic sleeve may augment any increase in LVEDP induced by the LV sleeve (Fig. 4F), while attenuating associated drops in EDV (Fig. 4G). In this work, we used two representative actuation volumes for the aortic sleeve and three volumes for the LV sleeve, which led to a total of eleven conditions being simulated; however, the volumes of each sleeve could be adjusted and modulated further to achieve additional hemodynamic states.

HFpEF represents a huge burden on the US healthcare system, with associated medical costs projected to exceed $25 million in 2030 (*53*). The efforts made towards the development of medical devices that can enhance cardiac mechanics in HFpEF patients have been largely hindered by the absence of adequate hemodynamic in vivo models, leading to the lack of FDA-approved device-based solutions for this condition. Most of the preclinical functional evaluation of HFpEF devices report testing on healthy in vivo models and unanimously emphasize the inability to simulate HFpEF hemodynamics as the primary limitation of their studies (*35, 42*). There is a broad variety of devices for HFpEF that are currently under development that would likely benefit from a reliable hemodynamic model of HFpEF. As an example of utility in acute evaluation of medical interventions for HFpEF, we demonstrated that our model could predict the hemodynamics associated with implant-free interatrial shunting and of an interatrial shunt device currently in clinical trials and seeking FDA approval (Fig. 5A-B). Although evaluating the efficacy of these interventions is not the purpose of this work, we showed that our model could recapitulate the hemodynamic changes associated with these treatment strategies, as predicted by computational models and human studies (*29, 33, 54, 55*). Particularly, we observed that shunting induced a shift of LV hemodynamics down the EDPVR curve due to a drop in preload (Fig. 5D), which led to a decrease in LVEDP (Fig. 5F), EDV (Fig. 5G), and SV (Fig. 5H). Further, diminished preload caused by shunting resulted in decreased systolic LVP (Fig. 5B) and dP_mean_ (Fig. 5E), likely governed by the Frank-Starling mechanism.

There are several advantages associated with this model. First, by relying on soft robotic actuatable sleeves, this model can recapitulate the hemodynamics of HFpEF in a controllable and highly tunable manner. The effect of each sleeve can be independently modulated by varying their respective actuation volume (or pressure). Having two degrees of freedom allows the models to mimic a wide variety of hemodynamic changes associated with HFpEF, from mild to severe filling dysfunction and pressure overload. As a result, our platform can capture the remarkably heterogeneous spectrum of HFpEF hemodynamics, which are observed clinically. This lends our model to medical device evaluation for a variety of hemodynamic subphenotypes of HFpEF and disease severities. Ultimately, this may lead to more reliable methods for device testing and the identification of subphenotype- or patient-specific device-based solutions for HFpEF. Finally, by acutely recreating the HFpEF hemodynamics, this model does not suffer from the prohibitive costs, lengthy development time, and limited throughput of previously reported large animal models of HFpEF induced by chronic pressure overload and/or metabolic disease.

One of the limitations of this approach involves its inability to predict changes in the long-term progression of HFpEF as affected by medical intervention. This is because this platform recapitulates the HFpEF hemodynamics using externally applied forces rather than biological processes. As a result, the effects of medical devices on remodeling processes and associated chronic hemodynamic changes cannot be comprehensively captured by this model. Secondly, although we demonstrated the ability to predict the acute hemodynamic effects of medical intervention by creating an LA-to-RA shunt, testing of extracardiac strategies, such as certain types of mechanical circulatory support strategies, may require some adjustments to the design of the LV sleeve, which may depend on the specific type and implantation approach of a given device. Future studies are warranted to further improve the clinical relevance of our proposed approach by exploring further the spectrum of medical devices for HFpEF that can be evaluated using this platform. Through partnership with industry, we also aim to conduct additional studies to comprehensively evaluate the efficacy of certain medical devices for specific hemodynamic phenotypes of HFpEF and severities of disease.

By describing a method that can recreate a broad spectrum of clinically relevant HFpEF hemodynamics in a highly controllable manner, this work can have a profound impact on current strategies for HFpEF device development and evaluation. This study could bridge the gap between in silico/in vitro testing and human trials for device-based solutions of HFpEF by overcoming the elevated costs and need for substantial resources for preclinical testing in a large animal model of disease, and by providing a better alternative to testing in healthy hemodynamic conditions, that is more reliable and tunable, hence more clinically relevant. We believe that this work represents a paradigm-shifting application of the use of soft robotics in in vivo disease modeling and device testing, which is a step forward towards addressing the lack of adequate treatment for HFpEF and alleviating its burden on healthcare and on the lives of the 32 million people currently suffering from this condition.

## MATERIALS AND METHODS

### Study design

This study involved the design of an implantable LV sleeve and the use of an aortic sleeve in a swine model (Yorkshire, male, ∼38-45 kg) for controllable recapitulation of HFpEF hemodynamics. The sleeves were sized using our cardiac magnetic resonance imaging database on swine within weight range and manufactured using materials and techniques detailed below. The sleeves were implanted in n = 3 swine and actuated using hydraulic pressure. The model hemodynamics were evaluated for i) two distinct actuation levels of the aortic sleeve (AoS), ii) three distinct levels of the LV sleeve (LVS), and iii) six combined levels, leading to a total of eleven hemodynamic scenarios. In each case, actuation was carried out at a constant infusion rate and held until stable hemodynamics were reached. On a subset of swine (n = 2), medical intervention was then simulated by percutaneous creation of a device-free interatrial shunt and implantation of an interatrial shunt device. Invasive hemodynamic monitoring was performed through LV and aortic catheterization for the entire duration of the studies. Metrics of pressure overload and diastolic dysfunction were obtained for model evaluation before and following intervention.

### Sleeve design and manufacturing

The LV sleeve was designed on SolidWorks (2019, Dassault Systèmes) using digital anatomies obtained from cardiac magnetic resonance images swine studies. The outer surface of the heart was offset by 10 mm to generate the sleeve geometry, which was then divided into four circumferential quadrants (each approximately 90 degrees apart), two for each ventricle. These quadrants were flattened to a plane to create the contours of the molds for manufacturing of the two TPU pockets for LV actuation. The design of the two molds was obtained by extrusion of the relative quadrant by the same offset (10 mm), and each mold was then 3D-printed using a rigid photopolymer (Veroblue, Stratasys) with an inkjet-based Objet30 3D-printer (Stratasys).

For each of the two LV molds, two TPU sheets (TPU, HTM 8001-M 80A shore polyether film, 0.012” thick, American Polyfilm) were vacuum-formed (Dental Vacuum Former, Yescom) to the shape of the molds. Each pair of TPU sheets was then heat-sealed at 320F for 8 seconds on a heat-press transfer machine using laser-cut negative acrylic molds to create enclosed and inflatable geometries. The edges of the two inflatable pockets were then heat-sealed to a 200-Denier TPU-coated fabric (Oxford fabric, Seattle fabrics Inc.), which was designed to be wrapped around the entire cardiac anatomy. Adjustable Velcro straps were added to the RV edges of the fabric to enable fastening. Two small openings were created through the fabric on one side of each of the two pockets to connect soft tubes (latex rubber 1/16“ ID 1/8” OD tubing, McMaster-Carr) as actuation lines through PVC connectors (polycarbonate plastic double-barbed tube fitting for 1/16“ tube ID, McMaster-Carr). The soft robotic aortic sleeve was designed and manufactured using a similar approach, previously described by our group (*51*).

### Surgical procedures and sleeve implantation

All animal procedures were approved by the Institutional Animal Care and Use Committee of our institute. In vivo studies were conducted on n = 3 Yorkshire swine housed in the Massachusetts Institute of Technology Department of Comparative Medicine Swine Facility.

After induction of anesthesia and endotracheal intubation, median sternotomy was performed using an oscillating saw. Dissection of the aortopulmonary window allowed for implantation of the aortic sleeve around the ascending aorta approximately 1-2 cm distal to the native aortic valve. The sleeve was tightened and secured using sutures. The LV sleeve was then fastened around the heart, taking care to orient the inflatable pockets around the LV, and tightened. The sleeve was secured with silk stay sutures to the pericardium to prevent downward translation during actuation.

Hemodynamic monitoring devices were placed using both endovascular and open-surgical access approaches. A vascular flow probe (ME 13 PXN, Transonic Systems Inc.) was placed around the descending thoracic aorta, and two PV catheters (Transonic Systems Inc.) were used to perform PV loop characterization with one 5F straight catheter inserted through femoral arterial access into the aortic arch and a 5F pigtail catheter placed transapically into the LV. Vital signs were monitored for the entire duration of the study.

### In vivo sleeve actuation

Actuation of the sleeves was performed using a syringe pump (70-3007 PHD ULTRA Syringe Pump Infuse/Withdraw, Harvard Apparatus). Three actuation volumes were used for the LV sleeve [mild (L1) = 40-45 mL, moderate (L2) = 50-55 mL, severe (L3) = 60-65 mL] and two for the aortic sleeve [moderate (A1) = 3.25-3.5 mL, severe (A2) = 4-4.25 mL]. The period of actuation was approximately 40 and 5 seconds for the LV and aortic sleeve, respectively. A stopcock was used to maintain a given actuation level of each sleeve throughout the duration of the study. The sleeves were actuated in isolation (A1-2, L1-L3) and in a combined fashion (A1L1, A1L2, A1L3, A2L1, A2L2, A2L3), leading to a total of 11 actuation modalities. The evaluation of interatrial shunting as an interventional approach for HFpEF was conducted on two representative cases, referred to as moderate (A1L2) and severe (A2L3).

### Interatrial shunting

Intervention was simulated in n = 2 swine. To this end, a device-free interatrial shunt was created percutaneously in one animal and a 6-mm AFR (Occlutech) was implanted in another animal. To create the interatrial shunt, ultrasound-guided access to the right femoral vein was obtained using the Seldinger technique. Under fluoroscopic guidance (OEC 9800), an 0.035” Bentson wire (Cook Medical) was then advanced into the RA. An 8.5Fr transseptal sheath (SJM Swartz, Abbott Laboratories) was then positioned in the RA and then across the interatrial septum into the LA. The wire was then placed into the left inferior pulmonary vein, and an 8-mm balloon (EverCross, Medtronic) was used to dilate the shunt. Once the atrial septal defect was created, deployment of the 6-mm AFR was performed under fluoroscopic guidance, using the manufacturer’s delivery system and pusher. Shunt patency was confirmed using epicardial color Doppler echocardiography (Philips EPiQ CVx with X5-1 transducer).

### Data acquisition

LV and aortic hemodynamics were monitored throughout the study following placement of the 5F PV catheters (Transonic, Inc.) in the LV and ascending arch, and the vascular flow probe on the descending thoracic aorta (Transonic, Inc.). The PV catheters were connected to the ADV500 PV system (Transonic Inc.) for measurements of LVP, LVV, and AoP. SV estimates of each animal were provided as inputs to the PV system for calibration, as obtained using trans-epicardial echocardiography (B-mode) on the EpiQ CVx system (Philips) prior to implantation of the sleeves. The vascular flow probe was connected to a two-channel flowmeter console (400 series, Transonic Inc.). Both the PV and flowmeter consoles were, in turn, connected to an eight-channel Powerlab system (ADInstruments) for real-time data acquisition with a 400-Hz sampling rate and recording. During the experiments, data were continuously monitored via LabChart software (Pro v8.1.16, ADInstruments). The default 50-Hz bandstop filter was applied to all signals. After the experiments, data were exported and processed in Matlab R2020a (MathWorks).

### Hemodynamic evaluation

Clinical metrics for the assessment of diastolic dysfunction and pressure overload were calculated for hemodynamic evaluation of the proposed model. The LVP_max_, LVEDP, EDV, and ESV were measured directly from the LVP and LVV tracings. Equations 1-3 were used for calculation of the SV, LVEF, and CO:

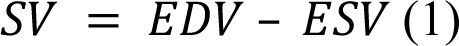

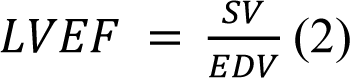

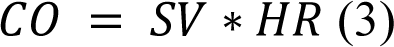

where HR indicates the heart rate in beats per minute (bpm).

For the assessment of pressure overload, dP_mean_ was calculated as the mean difference between the LVP and the AoP during cardiac ejection, each measured by the corresponding PV catheter.

The iEOA is calculated as in Equation 4, using the Gorlin equation (*56*):

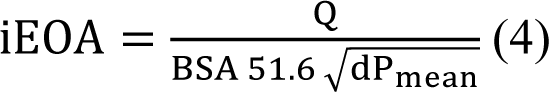

where *Q* is the flow through the aortic valve measured using the PV catheter in the LV and BSA indicates the body surface area of the animal.

The *Z_VA_* was measured using Equation 5(*57*):

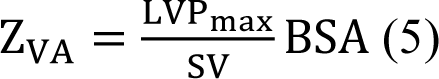

In Equations 4-5, the BSA (in m^2^) was estimated from the body weight (BW) (in kg) of the swine, using the Kelley formula (*58*) (Equation (6)):

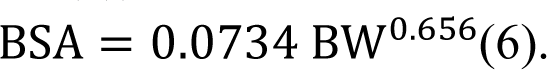

### Data analysis

All data were processed and analyzed on Matlab R2020a (Mathworks). Steady-state average and standard deviation values were obtained over ten consecutive heartbeats for each condition. Statistical significance between actuation levels was determined with respect to using two-tailed t-tests with a 95% confidence interval (P<0.05) on Matlab R2020a.

## One Sentence Summary

Implantable soft robotic sleeves can recapitulate the hemodynamics of HFpEF in a large animal model for device testing.

## Acknowledgments

We acknowledge Alison M Hayward from the Department of Comparative Medicine at Massachusetts Institute of Technology and Anna Spognardi from CBSET Inc. for their support with the in vivo studies.

## Funding

National Science Foundation 1847541 (ETR) NIH training grant T32 HL007734 (SXW)

Author contributions Conceptualization: LR, CO, SW, ETR Methodology: LR, CO, SW, MYS, ETR

Investigation: LR, CO, SW, DMQ, MYS, AM, ETR Visualization: LR, CO

Funding acquisition: ETR Project administration: ETR Supervision: ETR

Writing – original draft: LR

Writing – review & editing: LR, ETR

## Data and materials availability

All data are available in the main text.

## References

1. M. M. Kittleson, G. S. Panjrath, K. Amancherla, L. L. Davis, A. Deswal, D. L. Dixon, J. L. Januzzi, C. W. Yancy, 2023 ACC Expert Consensus Decision Pathway on Management of Heart Failure With Preserved Ejection Fraction: A Report of the American College of Cardiology Solution Set Oversight Committee., J. Am. Coll. Cardiol. 81, 1835–1878 (2023).

2. B. A. Borlaug, Evaluation and management of heart failure with preserved ejection fraction, Nat. Rev. Cardiol. 17, 559–573 (2020).

3. M. A. Pfeffer, A. M. Shah, B. A. Borlaug, Heart Failure with Preserved Ejection Fraction in Perspective*Circ*. Res. 124, 1598–1617 (2019).

4. M. A. Mamas, M. Sperrin, M. C. Watson, A. Coutts, K. Wilde, C. Burton, U. T. Kadam, C. S. Kwok, A. B. Clark, P. Murchie, I. Buchan, P. C. Hannaford, P. K. Myint, Do patients have worse outcomes in heart failure than in cancer? A primary care-based cohort study with 10-year follow-up in Scotland, Eur. J. Heart Fail. 19, 1095–1104 (2017).

5. M. M. Redfield, B. A. Borlaug, Heart Failure With Preserved Ejection Fraction, JAMA 329, 827 (2023).

6. S. M. Dunlay, V. L. Roger, M. M. Redfield, Epidemiology of heart failure with preserved ejection fraction*Nat*. Rev. Cardiol. 14, 591–602 (2017).

7. C. S. P. Lam, A. A. Voors, R. A. De Boer, S. D. Solomon, D. J. Van Veldhuisen, Heart failure with preserved ejection fraction: From mechanisms to therapies*Eur*. Heart J. (2018), doi:10.1093/eurheartj/ehy301.

8. B. A. Borlaug, W. J. Paulus, Heart failure with preserved ejection fraction: pathophysiology, diagnosis, and treatment, Eur. Heart J. 32, 670–679 (2011).

9. A. Ferron, F. Francisqueti, I. Minatel, C. Silva, S. Bazan, K. Kitawara, J. Garcia, C. Corrêa, F. Moreto, A. Ferreira, Association between Cardiac Remodeling and Metabolic Alteration in an Experimental Model of Obesity Induced by Western Diet, Nutrients 10, 1675 (2018).

10. W. J. Paulus, M. R. Zile, From Systemic Inflammation to Myocardial Fibrosis, Circ. Res. 128, 1451–1467 (2021).

11. K. Modesto, P. P. Sengupta, Myocardial Mechanics in Cardiomyopathies, Prog. Cardiovasc. Dis. 57, 111–124 (2014).

12. S. Ihne, C. Morbach, L. Obici, G. Palladini, S. Störk, Amyloidosis in Heart Failure, Curr. Heart Fail. Rep. 16, 285–303 (2019).

13. E. Braunwald, Cardiomyopathies, Circ. Res. 121, 711–721 (2017).

14. M. R. Dweck, S. Joshi, T. Murigu, A. Gulati, F. Alpendurada, A. Jabbour, A. Maceira, I. Roussin, D. B. Northridge, P. J. Kilner, S. A. Cook, N. A. Boon, J. Pepper, R. H. Mohiaddin, D. E. Newby, D. J. Pennell, S. K. Prasad, Left ventricular remodeling and hypertrophy in patients with aortic stenosis: insights from cardiovascular magnetic resonance, J. Cardiovasc. Magn. Reson. 14, 50 (2012).

15. C. Tsioufis, G. Georgiopoulos, D. Oikonomou, C. Thomopoulos, N. Katsiki, A. Kasiakogias, C. Chrysochoou, D. Konstantinidis, T. Kalos, D. Tousoulis, Hypertension and Heart Failure with Preserved Ejection Fraction: Connecting the Dots, Curr. Vasc. Pharmacol. 16 (2017), doi:10.2174/1570161115666170414120532.

16. B. A. Carabello, W. J. Paulus, Aortic stenosis, Lancet 373, 956–966 (2009).

17. M. Penicka, J. Bartunek, H. Trakalova, H. Hrabakova, M. Maruskova, J. Karasek, V. Kocka, Heart Failure With Preserved Ejection Fraction in Outpatients With Unexplained Dyspnea. A Pressure-Volume Loop Analysis, J. Am. Coll. Cardiol. 55, 1701–1710 (2010).

18. P. A. Heidenreich, B. Bozkurt, D. Aguilar, L. A. Allen, J. J. Byun, M. M. Colvin, A. Deswal, M. H. Drazner, S. M. Dunlay, L. R. Evers, J. C. Fang, S. E. Fedson, G. C. Fonarow, S. S. Hayek, A. F. Hernandez, P. Khazanie, M. M. Kittleson, C. S. Lee, M. S. Link, C. A. Milano, L. C. Nnacheta, A. T. Sandhu, L. W. Stevenson, O. Vardeny, A. R. Vest, C. W. Yancy, 2022 AHA/ACC/HFSA Guideline for the Management of Heart Failure: A Report of the American College of Cardiology/American Heart Association Joint Committee on Clinical Practice Guidelines, Circulation 145, e895–e1032 (2022).

19. L. Fauchier, A. Bisson, A. Bodin, Heart failure with preserved ejection fraction and atrial fibrillation: recent advances and open questions, BMC Med. 21, 54 (2023).

20. M. M. Hoeper, C. S. P. Lam, J.-L. Vachiery, J. Bauersachs, C. Gerges, I. M. Lang, D. Bonderman, K. M. Olsson, J. S. R. Gibbs, P. Dorfmuller, M. Guazzi, N. Galiè, A. Manes, M. L. Handoko, A. Vonk-Noordegraaf, M. Lankeit, S. Konstantinides, R. Wachter, C. Opitz, S. Rosenkranz, Pulmonary hypertension in heart failure with preserved ejection fraction: a plea for proper phenotyping and further research†, *Eur. Heart J.*, ehw597 (2016).

21. F. Berglund, P. Piña, C. J. Herrera, Right ventricle in heart failure with preserved ejection fraction, Heart 106, 1798–1804 (2020).

22. V. G. Florea, J. N. Cohn, The autonomic nervous system and heart failure*Circ*. Res. 114, 1815–1826 (2014).

23. Y. Xi, J. Cheng, Dysfunction of the autonomic nervous system in atrial fibrillation*J*. Thorac. Dis. 7, 193–198 (2015).

24. Y. Lytvyn, P. Bjornstad, J. A. Udell, J. A. Lovshin, D. Z. I. Cherney, Sodium Glucose Cotransporter-2 Inhibition in Heart Failure, Circulation 136, 1643–1658 (2017).

25. Y. Xie, Y. Wei, D. Li, J. Pu, H. Ding, X. Zhang, Mechanisms of SGLT2 Inhibitors in Heart Failure and Their Clinical Value, J. Cardiovasc. Pharmacol. 81, 4–14 (2023).

26. A. Pandey, A. Parashar, D. J. Kumbhani, S. Agarwal, J. Garg, D. Kitzman, B. D. Levine, M. Drazner, J. D. Berry, Exercise Training in Patients With Heart Failure and Preserved Ejection Fraction, Circ. Hear. Fail. 8, 33–40 (2015).

27. A. Jasinska-Piadlo, P. Campbell, Management of patients with heart failure and preserved ejection fraction, Heart 109, 874–883 (2023).

28. L. Rosalia, C. Ozturk, S. Shoar, Y. Fan, G. Malone, F. H. Cheema, C. Conway, R. A. Byrne, G. P. Duffy, A. Malone, E. T. Roche, A. Hameed, Device-Based Solutions to Improve Cardiac Physiology and Hemodynamics in Heart Failure With Preserved Ejection Fraction, JACC Basic to Transl. Sci. 6, 772–795 (2021).

29. M. Granegger, C. Gross, D. Siemer, A. Escher, S. Sandner, M. Schweiger, G. Laufer, D. Zimpfer, Comparison of device-based therapy options for heart failure with preserved ejection fraction: a simulation study, Sci. Rep. 12, 5761 (2022).

30. N. Berry, L. Mauri, T. Feldman, J. Komtebedde, D. J. van Veldhuisen, S. D. Solomon, J. M. Massaro, S. J. Shah, Transcatheter InterAtrial Shunt Device for the treatment of heart failure: Rationale and design of the pivotal randomized trial to REDUCE Elevated Left Atrial Pressure in Patients with Heart Failure II (REDUCE LAP-HF II): Rationale and design of REDUCE LA (2020).

31. J. Rodés-Cabau, M. Bernier, I. J. Amat-Santos, T. Ben Gal, L. Nombela-Franco, B. García del Blanco, A. Kerner, S. Bergeron, M. del Trigo, P. Pibarot, S. Shkurovich, N. Eigler, W. T. Abraham, Interatrial Shunting for Heart Failure: Early and Late Results From the First-in-Human Experience With the V-Wave System, JACC Cardiovasc. Interv. 11, 2300–2310 (2018).

32. C. Paitazoglou, M. W. Bergmann, R. Özdemir, R. Pfister, J. Bartunek, T. Kilic, A. Lauten, A. Schmeisser, M. Zoghi, S. Anker, H. Sievert, F. Mahfoud, One-year results of the first-in-man study investigating the Atrial-Flow-Regulator for left-atrial shunting in symptomatic heart failure patients: the PRELIEVE study, Eur. J. Heart Fail. (2021), doi:10.1002/ejhf.2119.

33. J. E. Udelson, C. M. Barker, G. Wilkins, B. Wilkins, R. Gooley, S. Lockwood, B. J. Potter, C. U. Meduri, P. S. Fail, D. J. Solet, K. Feldt, J. M. Kriegel, T. Shaburishvili, No-Implant Interatrial Shunt for HFpEF, JACC Hear. Fail. (2023), doi:10.1016/j.jchf.2023.01.024.

34. T. Simard, M. Labinaz, F. Zahr, B. Nazer, W. Gray, J. Hermiller, S. P. Chaudhry, L. Guimaraes, F. Philippon, P. Eckman, J. Rodés-Cabau, P. Sorajja, B. Hibbert, Percutaneous Atriotomy for Levoatrial–to–Coronary Sinus Shunting in Symptomatic Heart Failure: First-in-Human Experience, JACC Cardiovasc. Interv. (2020), doi:10.1016/j.jcin.2020.02.022.

35. Y. Feld, S. Dubi, Y. Reisner, E. Schwammenthal, R. Shofti, A. Pinhasi, S. Carasso, A. Elami, Energy transfer from systole to diastole: A novel device-based approach for the treatment of diastolic heart failure, Acute Card. Care (2011), doi:10.3109/17482941.2011.634012.

36. C. Ozturk, L. Rosalia, E. T. Roche, A Multi-Domain Simulation Study of a Pulsatile-Flow Pump Device for Heart Failure With Preserved Ejection Fraction, Front. Physiol. 13 (2022), doi:10.3389/fphys.2022.815787.

37. C. Miyagi, J. H. Karimov, B. D. Kuban, T. Miyamoto, S. Sale, C. Flick, R. C. Starling, K. Fukamachi, Development of the Left Atrial Assist Device for Patients with Heart Failure with Preserved Ejection Fraction: First In Vivo Results, *J. Hear*. Lung Transplant. 40, S176–S177 (2021).

38. A. Escher, Y. Choi, F. Callaghan, B. Thamsen, U. Kertzscher, M. Schweiger, M. Hübler, M. Granegger, A Valveless Pulsatile Pump for Heart Failure with Preserved Ejection Fraction: Hemo- and Fluid Dynamic Feasibility, Ann. Biomed. Eng. 48, 1821–1836 (2020).

39. E. Gude, A. E. Fiane, Can mechanical circulatory support be an effective treatment for HFpEF patients?, Hear. Fail. Rev. 2021 1, 1–9 (2021).

40. A. Rosenzweig, The Growing Importance of Basic Models of Cardiovascular Disease, Circ. Res. 130, 1743–1746 (2022).

41. C. Miyagi, T. Miyamoto, T. Kuroda, J. H. Karimov, R. C. Starling, K. Fukamachi, Large animal models of heart failure with preserved ejection fraction, Hear. Fail. Rev. 2021, 1–14 (2021).

42. Y. Feld, in PCR innovators day, (PCR, 2019).

43. W. M. Yarbrough, R. Mukherjee, R. E. Stroud, W. T. Rivers, J. M. Oelsen, J. A. Dixon, S. R. Eckhouse, J. S. Ikonomidis, M. R. Zile, F. G. Spinale, Progressive induction of left ventricular pressure overload in a large animal model elicits myocardial remodeling and a unique matrix signature, J. Thorac. Cardiovasc. Surg. 143, 215–223 (2012).

44. M. Gyöngyösi, N. Pavo, D. Lukovic, K. Zlabinger, A. Spannbauer, D. Traxler, G. Goliasch, L. Mandic, J. Bergler-Klein, A. Gugerell, A. Jakab, Z. Szankai, L. Toth, R. Garamvölgyi, G. Maurer, F. Jaisser, F. Zannad, T. Thum, S. Bátkai, J. Winkler, Porcine model of progressive cardiac hypertrophy and fibrosis with secondary postcapillary pulmonary hypertension, J. Transl. Med. 15, 202 (2017).

45. W. M. Torres, S. C. Barlow, A. Moore, L. A. Freeburg, A. Hoenes, H. Doviak, M. R. Zile, T. Shazly, F. G. Spinale, Changes in Myocardial Microstructure and Mechanics With Progressive Left Ventricular Pressure Overload, JACC Basic to Transl. Sci. 5, 463–480 (2020).

46. V. K. Munagala, C. Y. T. Hart, J. C. Burnett, D. M. Meyer, M. M. Redfield, Ventricular Structure and Function in Aged Dogs With Renal Hypertension, Circulation 111, 1128–1135 (2005).

47. O. Sorop, I. Heinonen, M. van Kranenburg, J. van de Wouw, V. J. de Beer, I. T. N. Nguyen, Y. Octavia, R. W. B. van Duin, K. Stam, R.-J. van Geuns, P. A. Wielopolski, G. P. Krestin, A. H. van den Meiracker, R. Verjans, M. van Bilsen, A. H. J. Danser, W. J. Paulus, C. Cheng, W. A. Linke, J. A. Joles, M. C. Verhaar, J. van der Velden, D. Merkus, D. J. Duncker, Multiple common comorbidities produce left ventricular diastolic dysfunction associated with coronary microvascular dysfunction, oxidative stress, and myocardial stiffening, Cardiovasc. Res. 114, 954–964 (2018).

48. C. Mühlfeld, A. Rajces, M. Manninger, A. Alogna, M. Wierich, D. Scherr, H. Post, J. Schipke, A transmural gradient of myocardial remodeling in early-stage heart failure with preserved ejection fraction in the pig, J. Anat. 236, 531–539 (2020).

49. T. D. Olver, J. C. Edwards, T. J. Jurrissen, A. B. Veteto, J. L. Jones, C. Gao, C. Rau, C. M. Warren, P. J. Klutho, L. Alex, S. C. Ferreira-Nichols, J. R. Ivey, P. K. Thorne, K. S. McDonald, M. Krenz, C. P. Baines, R. J. Solaro, Y. Wang, D. A. Ford, T. L. Domeier, J. Padilla, R. S. Rector, C. A. Emter, Western Diet-Fed, Aortic-Banded Ossabaw Swine: A Preclinical Model of Cardio-Metabolic Heart Failure, JACC Basic to Transl. Sci. 4, 404–421 (2019).

50. L. Rosalia, C. Ozturk, D. Goswami, J. Bonnemain, S. X. Wang, B. Bonner, J. C. Weaver, R. Puri, S. Kapadia, C. T. Nguyen, E. T. Roche, Soft robotic patient-specific hydrodynamic model of aortic stenosis and ventricular remodeling, *Sci*. Robot. 8 (2023), doi:10.1126/scirobotics.ade2184.

51. L. Rosalia, C. Ozturk, J. Coll-Font, Y. Fan, Y. Nagata, M. Singh, D. Goswami, A. Mauskapf, S. Chen, R. A. Eder, E. M. Goffer, J. H. Kim, S. Yurista, B. P. Bonner, A. N. Foster, R. A. Levine, E. R. Edelman, M. Panagia, J. L. Guerrero, E. T. Roche, C. T. Nguyen, A soft robotic sleeve mimicking the haemodynamics and biomechanics of left ventricular pressure overload and aortic stenosis, *Nat*. Biomed. Eng. 6, 1134–1147 (2022).

52. T. A. McDonagh, M. Metra, M. Adamo, R. S. Gardner, A. Baumbach, M. Böhm, H. Burri, J. Butler, J. Čelutkienė, O. Chioncel, J. G. F. Cleland, A. J. S. Coats, M. G. Crespo-Leiro, D. Farmakis, M. Gilard, S. Heymans, A. W. Hoes, T. Jaarsma, E. A. Jankowska, M. Lainscak, C. S. P. Lam, A. R. Lyon, J. J. V McMurray, A. Mebazaa, R. Mindham, C. Muneretto, M. Francesco Piepoli, S. Price, G. M. C. Rosano, F. Ruschitzka, A. Kathrine Skibelund, R. A. de Boer, P. Christian Schulze, M. Abdelhamid, V. Aboyans, S. Adamopoulos, S. D. Anker, E. Arbelo, R. Asteggiano, J. Bauersachs, A. Bayes-Genis, M. A. Borger, W. Budts, M. Cikes, K. Damman, V. Delgado, P. Dendale, P. Dilaveris, H. Drexel, J. Ezekowitz, V. Falk, L. Fauchier, G. Filippatos, A. Fraser, N. Frey, C. P. Gale, F. Gustafsson, J. Harris, B. Iung, S. Janssens, M. Jessup, A. Konradi, D. Kotecha, E. Lambrinou, P. Lancellotti, U. Landmesser, C. Leclercq, B. S. Lewis, F. Leyva, A. Linhart, M.-L. Løchen, L. H. Lund, D. Mancini, J. Masip, D. Milicic, C. Mueller, H. Nef, J.-C. Nielsen, L. Neubeck, M. Noutsias, S. E. Petersen, A. Sonia Petronio, P. Ponikowski, E. Prescott, A. Rakisheva, D. J. Richter, E. Schlyakhto, P. Seferovic, M. Senni, M. Sitges, M. Sousa-Uva, C. G. Tocchetti, R. M. Touyz, C. Tschoepe, J. Waltenberger, M. Adamo, A. Baumbach, M. Böhm, H. Burri, J. Čelutkienė, O. Chioncel, J. G. F. Cleland, A. J. S. Coats, M. G. Crespo-Leiro, D. Farmakis, R. S. Gardner, M. Gilard, S. Heymans, A. W. Hoes, T. Jaarsma, E. A. Jankowska, M. Lainscak, C. S. P. Lam, A. R. Lyon, J. J. V McMurray, A. Mebazaa, R. Mindham, C. Muneretto, M. F. Piepoli, S. Price, G. M. C. Rosano, F. Ruschitzka, A. K. Skibelund, 2021 ESC Guidelines for the diagnosis and treatment of acute and chronic heart failure, Eur. Heart J. 42, 3599–3726 (2021).

53. P. A. Heidenreich, N. M. Albert, L. A. Allen, D. A. Bluemke, J. Butler, G. C. Fonarow, J. S. Ikonomidis, O. Khavjou, M. A. Konstam, T. M. Maddox, G. Nichol, M. Pham, I. L. Piña, J. G. Trogdon, Forecasting the impact of heart failure in the united states a policy statement from the american heart association, Circ. Hear. Fail. 6, 606–619 (2013).

54. M. B. Patel, B. P. Samuel, R. E. Girgis, M. A. Parlmer, J. J. Vettukattil, Implantable atrial flow regulator for severe, irreversible pulmonary arterial hypertension, EuroIntervention 11, 706–709 (2015).

55. R. Rajeshkumar, S. Pavithran, K. Sivakumar, J. J. Vettukattil, Atrial septostomy with a predefined diameter using a novel occlutech atrial flow regulator improves symptoms and cardiac index in patients with severe pulmonary arterial hypertension, Catheter. Cardiovasc. Interv. 90, 1145–1153 (2017).

56. N. Saikrishnan, G. Kumar, F. J. Sawaya, S. Lerakis, A. P. Yoganathan, Accurate Assessment of Aortic Stenosis, Circulation 129, 244–253 (2014).

57. R.-J. Nuis, J. A. Goudzwaard, M. J. A. G. de Ronde-Tillmans, H. Kroon, J. F. Ooms, M. P. van Wiechen, M. L. Geleijnse, F. Zijlstra, J. Daemen, N. M. Van Mieghem, F. U. S. Mattace-Raso, M. J. Lenzen, P. P. T. de Jaegere, Impact of Valvulo-Arterial Impedance on Long-Term Quality of Life and Exercise Performance After Transcatheter Aortic Valve Replacement, Circ. Cardiovasc. Interv. 13 (2020), doi:10.1161/CIRCINTERVENTIONS.119.008372.

58. T. Itoh, M. Kawabe, T. Nagase, H. Matsushita, M. Kato, M. Miyoshi, K. Miyahara, Body surface area measurement in juvenile miniature pigs using a computed tomography scanner, Exp. Anim. 66, 229–233 (2017).

